# History-dependent Biases in Perceptual Decisions Depend on NMDA Receptors

**DOI:** 10.64898/2026.01.12.699039

**Authors:** Alessandro Toso, Ayelet Arazi, Anke Braun, Rafael Marin, Philipp Sterzer, Jaime de la Rocha, Tobias H. Donner

## Abstract

Perceptual decisions are governed not only by current sensory input but also by the history of previous stimuli, choices, or outcomes. Such history-dependent biases evolve across multiple timescales ranging from seconds to minutes. Synaptic plasticity is one possible mechanism mediating such effects. The by N-methyl-aspartate (NMDA) glutamate receptor is critical for synaptic plasticity, and indirect evidence points to its role in the generation of history-dependent biases. Here, we pharmacologically blocked NMDA receptors in healthy human participants performing a two-alternative visual decision-making task. Participants’ choices were biased by the accumulation of fluctuating sensory evidence within each trial as well as the accumulation of across-trial sequences of stimuli, which enabled them to adapt to changing stimulus statistics across blocks. NMDA receptor blockade reduced the task-adaptive across trials, history-dependent biases. In contrast, enhancing GABA-A receptor activity in the same participants increased the transiency of the within-trial evidence accumulation dynamics. Our findings provide direct evidence for the importance of NMDA receptor function in history-dependent biases in the human brain.

## Introduction

Decisions are shaped not only by the current sensory input, but also by prior knowledge and expectations. For example, during foraging behaviour, choices are governed by the history of previous actions and experienced rewards^1,2^. Even during simple perceptual judgments, choices are not only based on the current sensory input, but also strongly biased by the trial history. Such history-dependent biases in perceptual decision-making are prominent in both humans^3–7^ and animals^8–14^. Within frameworks conceptualizing a perceptual judgement as a combination of incoming evidence with prior expectations^15^, history-dependent biases can be understood as the behavioral expression of dynamic prior expectations that are shaped by past choices and stimuli ^10,12,16^. The states of natural sensory environments have rich temporal structure and are commonly stable over prolonged timescales compared to the rapid fluctuations of the raw sensory input^17^. To leverage this temporal structure, the brain adapts the construction of history-dependent decision biases to sensory environment^7,12^.

The mechanisms underlying such history-dependent biases have remained debated. These biases can unfold over multiple timescales ranging from a few seconds^16^ to tens of seconds^6,7^ to minutes^18,19^ - in other words, substantially longer than the timescales that individual neurons are equipped with implying that choice history biases are an emergent network property^20^. Indeed, the timescales of history biases may even be longer than those of persistent, stimulus- or choice-selective cortical activity that is often observed during tasks requiring short-term memory and/or the accumulation of sensory evidence^21,22^. For these reasons, changes in synaptic efficacy have been implicated in carrying such selective information over trials^23,24,72^.

The N-methyl-aspartate (NMDA) receptor is critical for long-term synaptic plasticity^25^ and has been hypothesized to also play a role in plasticity on the intermediate (seconds to tens of seconds) timescales that shape sequential effects in decision-making^24,26,27^. History-dependent biases in working memory tasks are reduced in patients with putative NMDA receptor hypo-function - due to anti-NMDA encephalitis^24^ or (early stage) schizophrenia^24,28,29^. However, these patients may have alterations in brain circuits other than just NMDA hypofunction^30^. Further, these clinical populations have so far only been tested in tasks other than perceptual categorization tasks, in which adaptive choice history biases have been established in healthy humans and rodents, which may rely on different mechanisms^7,12,16,31^. Finally, theoretical work also implicates NMDA receptors in the accumulation of sensory evidence occurring in the presence of sensory input^32,33^. Direct evidence from manipulations of NMDA receptors in healthy individuals, in a task entailing both, within-trial accumulation of evidence as well as across-trial history biases, is needed to understand the role of NMDA receptors in the multi-timescale dynamics of perceptual decision-making.

Here, we combined a two-alternative visual evidence accumulation task with placebo-controlled, low dose pharmacological blockade of NMDA receptors. To test for neurochemical specificity, we also (in the same participants) manipulated inhibitory GABA-A receptors, which also play a critical role in neural circuit models of perceptual decision-making^20^. This experimental design enabled us to test how both key neurotransmitter receptors shape perceptual decision formation across different timescales. We found distinct effects of both manipulations: As expected, NMDA-blockade reduced across-trial history-dependent biases, while GABA-A boost altered the dynamics of within-trial evidence accumulation.

## Results

### Adjusting trial-history biases to environmental statistics

To characterize the mechanisms underlying the generation of trial-history biases, twenty healthy human participants performed a visual categorization task under placebo-controlled pharmacological interventions (Figure 1). Each participant performed the task across six experimental sessions, each preceded by the oral administration of the NMDA receptor blocker memantine (15 mg), GABA-A receptor agonist lorazepam (1 mg), or a placebo in a double-blind, randomized, crossover design (Figure S1, see Methods for details). Because NMDA and GABA-A receptors have both been implicated in the cortical circuit mechanisms of decision formation^21,34^, the GABA-A manipulation served as a test for specificity of any NMDA effects. Both drugs were administered at low dosage to minimize non-specific side effects.

**Figure 1.**
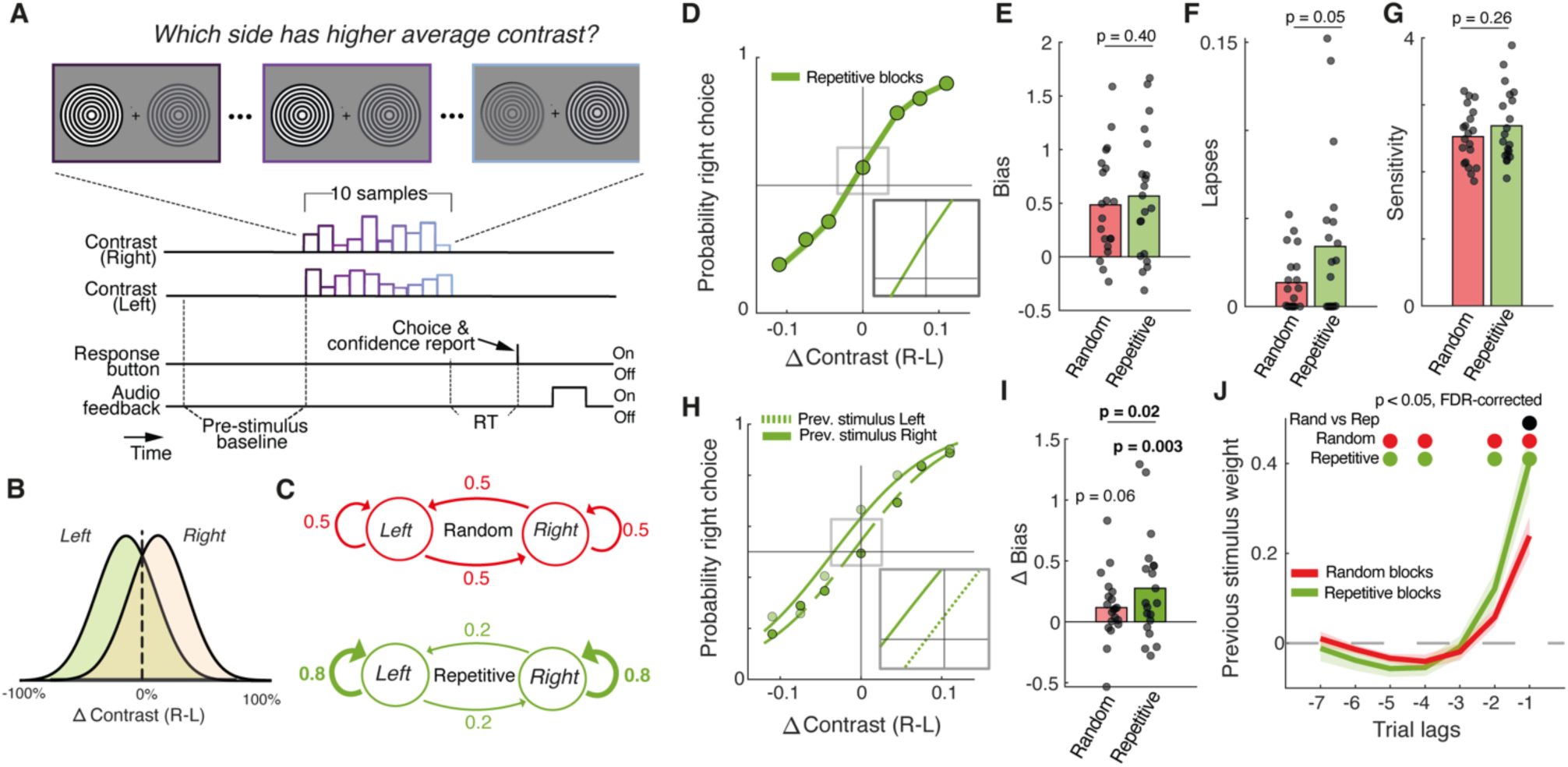
Behavioral task and trial history bias. **(A)** Schematic of task events during an example trial. A sequence of fluctuating contrast samples is shown in each visual hemifield. After stimulus offset, participants report the side with the larger mean contrast (left versus right) combined with their confidence in the correctness of that judgment (high versus low) by pushing one of four buttons. See main text for details. Auditory feedback was provided after responses. **(B)** Stimulus categories *L*eft and *Right* ared efined by the difference in their generative mean of the two contrast distributions (see main text). **(C)** Task environments are defined by the probability to repeat the previous stimulus category: in the Random environment the repetition probability was 0.5 whereas in the Repetitive the repetition probability was 0.8. **(D)** Probability of ‘right’ choice as a function of signed evidence strength (see main text) computed for repetitive environment. Data points, group average. Line, psychometric function fit. Inset, close-up of curve. **(E-G)** Psychometric fitted parameters bias (E), lapse rate (F), and sensitivity (G). Data points, individual participants; bars, group average. P-values obtained from two-sided permutation test. **(H)** As panel (D), but conditioned by previous stimulus category (*Left* or *Right*). We see a clear horizontal shift in opposite directions according to the previous stimulus side. **(I)** Difference in bias parameter between previous stimulus categories (horizontal shift between dashed lines and solid lines in H), separately for the two environments. Positive values indicate the tendency to repeat the previous stimulus category. Data points, individual participants; bars, group average. P-values (two-sided permutation test). **(J)** History kernels quantifying the impact of previous stimulus categories as a function of trial lag on current choice. Panels D-J show data from placebo sessions only. Colors indicate the two sensory environments. Lines, group average, shaded area, SEM. Upper marks, p < 0.05; FDR-corrected.

In the task, participants were asked to judge on each trial, which of two visual fluctuating stimuli presented in the left and right visual hemifields had a stronger mean contrast (Figure 1A; Methods). Each stimulus consisted of a sequence of ten successive circular gratings (100 ms duration each). The contrast of each grating was randomly sampled from a Gaussian distribution (standard deviation: 15% Michelson contrast) which had a mean slightly above 50% on one side, and slightly below 50% on the other side, respectively, titrated to keep performance around 75% correct (Methods). The difference between both means, called “delta contrast” was the signed evidence strength. Its sign determined the two stimulus categories, abbreviated as *Right* (delta > 0) and *Left* (delta < 0) (Figure 1B), and its absolute value determined decision difficulty. Participants reported the inferred category by pressing a button with the index finger of the right or the left hand, respectively. Auditory feedback about the correctness of participants’ choices was provided after responses.

To promote the integration of decision evidence across trials, we introduced two environments that varied across the session in trial blocks. Each environment was defined by the statistics of the across-trial sequence of stimulus categories (*Left* or *Right*) and was either Random (probability of repeating the previous category *p(repeat)* = 0.5) or Repetitive (repetition probability *p_r_* = 0.8; Methods). Participants were not informed about the existence and structure of these two environments. The Repetitive environment was specifically introduced to induce consistent trial history-dependent biases based on previous work using similar designs^7,12,16^ because history biases in random environments exhibit considerable inter-individual differences^35^. We first present analyses of participants’ behavior under placebo, followed by a quantification of the effects of both drugs.

### Adaptive bias dependent on stimulus history

Choice probability lawfully scaled with the signed evidence strength in both environments (Figure 1D). Fits of the corresponding psychometric function (logistic regression, Methods) provided no evidence for a difference between overall bias (Figure 1E), in the two environments. The overall bias was towards the right choice in the majority of individuals, possibly reflecting participants’ handedness (Methods). The environments, likewise, did not yield any differences in stimulus-independent errors (lapse rate, Figure 1F), or sensitivity (slope, Figure 1G).

As in previous work using the same statistical structure in the context of a random dot motion task^16^, choices were biased by trial history in a manner that reflected the correlation structure of the environment (Figure 1H,I). We used the same logistic regression model mentioned above for the psychometric function to quantify these history-dependent biases (^7,16,36^, Methods) and decomposed these biases into the impact of previous stimulus categories (Figure 1J) and choices (Figure S2) from the last seven trials on the current choice, respectively. This revealed consistently positive contributions (i.e., beta weights) for the two previously presented stimulus categories, combined with a negative contribution of the two previous choices indicative of a “loose-switch” strategy^36^ (Figure S2). This strategy was likely governed by the immediate feedback about choice outcome that participants received after each trial (Figure 1A).

Critically, the previous stimulus weights adapted to the statistical structure, being larger for the Repetitive environment (Figure 1J), an effect not observed for previous choice (Figure S2) and consistent with previous work quantifying history biases in perceptual choice task of similar structure^16^. Because the feedback disambiguated the previous stimulus category even when the corresponding evidence happened to be weak, participants could always use their knowledge about the previous category to build a prior expectation for the next decision in the Repetitive environment^16^.

The model used above was purely statistical and did not describe any mechanism underlying participants’ choice behavior. The behavior, and in particular the history-dependent biases, were also well fit by a generative model adapted from previous rodent work^12^ (Figure S3). Different from the statistical model, the generative model was able to track the statistics of the stimulus sequence and develop a choice bias adapted to each environment. Consequently, it could be fit to all data without sorting by sensory environment. In the model, not only choices and stimulus categories, but also transitions (i.e., repetitions or alternations) are accumulated across trials (Figure S3A; Methods), which enables the model to learn the environmental context (i.e., Repetitive versus Random). Critically, also this model uncovered a positive and significant contribution of previous stimulus category (Figure S3E) accompanied by a negative contribution of previous choices (Figure S3F), mirroring the results of the statistical model. Further, the fits showed a positive so-called transition evidence (Figure S3H), which captures participants’ ongoing learning of the stimulus statistics (transition probabilities) that produced the history biases adjustment to the different environments evident in Figure 1I-J.

In sum, participants exhibited a consistent bias towards choosing the side corresponding to the previously presented stimulus category, particularly pronounced in the Repetitive sensory environment. This tendency was evident in two independent modeling approaches and in line with normative theory^37^ as well as empirical findings from different species^12,16^. We next tested if this bias component was affected by the manipulation of NMDA receptors.

### NMDA-R blockade reduces stimulus history-dependent bias

As expected from the low dosage of the drugs, both memantine and lorazepam had little impact on current-trial performance (Figure 2A-D). We found no statistically significant effects on lapse rates that captured stimulus-independent errors (Figure 2B), sensitivity (Figure 2C), nor overall bias (Figure 2D). The fact that evidence sensitivity did not decrease and stimulus-independent errors did not increase under either drug compared to placebo verifies the intended subtlety of the manipulations and avoids interpretational complications due to lowered arousal under the drugs.

**Figure 2.**
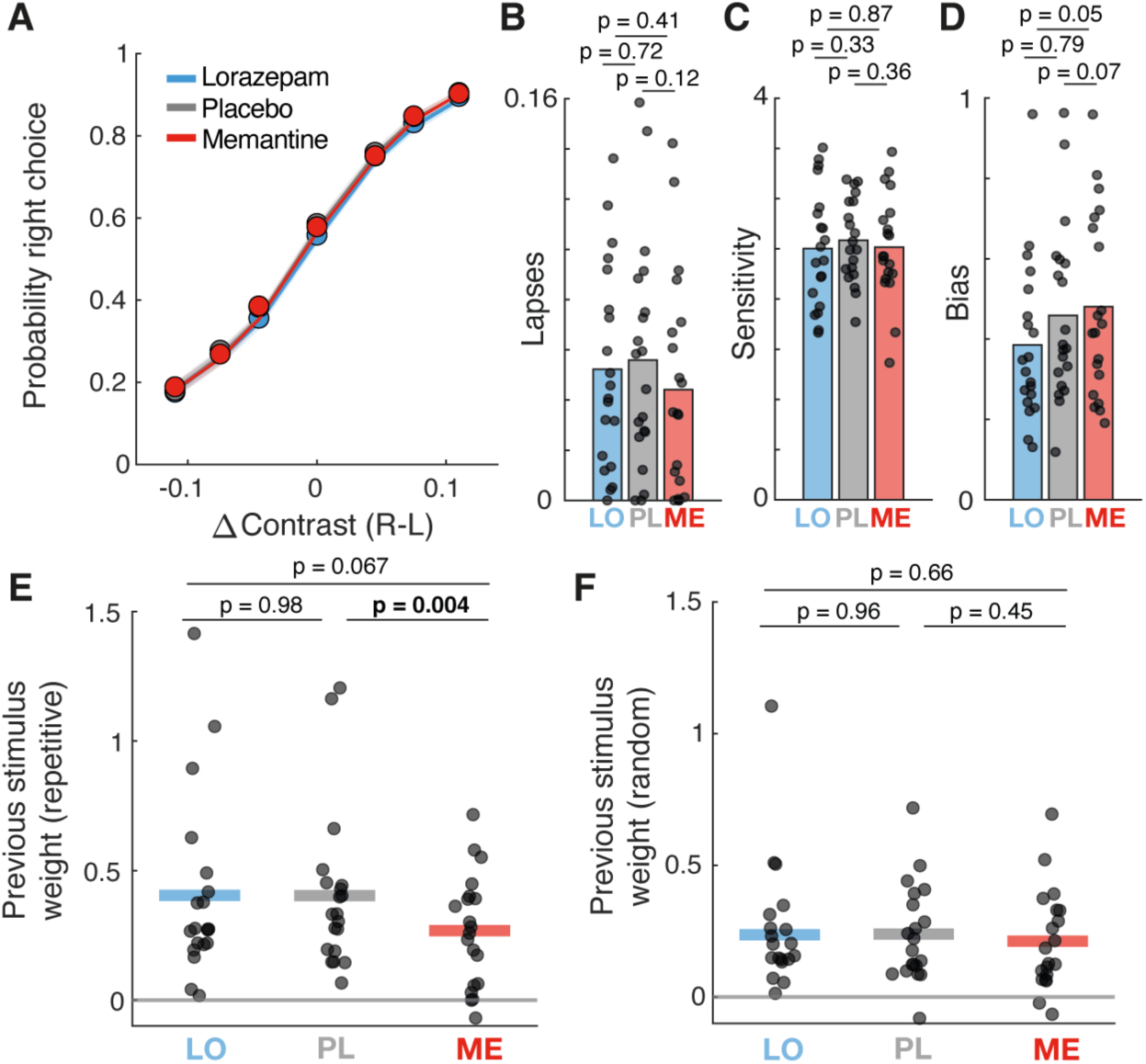
Drug-induced changes in stimulus history bias. Panels show the parameters quantifying choice behavior split by pharmacological condition. **(A)** Probability of right-side choice as a function of signed evidence strength. Data points, group average. Lines, psychometric function fits. **(B)** Lapse rates. **(C)** Sensitivity. **(D)** Overall bias. **(E)** Impact of previous stimulus category in repetitive environment. **(F)** Same as E for Random P-values in all panels are from two-sided permutation tests. Horizontal lines, group average; data points, individual participants.

Critically, we found that the NMDA blockade through memantine strongly and highly significantly reduced the stimulus-dependent history bias, compared to both, lorazepam and placebo in repetitive environment only (Figure 2E). This effect was highly specific: it was unrelated to the memantine-induced effect on lapse rate (Figure S4A) and was absent for previous choice (Figure S4B-C) as well as for the Random environment (Figure 2F). In the next section, we will show that memantine, at this low dosage also did not affect the accumulation of the fluctuating stimulus evidence within trials.

### NMDA-R blockade does not affect within-trial accumulation of evidence

In current cortical circuit models^34,38^, the integration of sensory evidence towards categorical decisions unfolding within trials^39,40^ emerges from recurrent excitation (via NMDA receptors) within, and competition (via GABA-A receptors) between two pools of excitatory neurons integrating evidence for one or the other choice^20,34^. In such circuit models, changing the conductance of NMDA (or GABA-A) synapses alters the temporal profile of the weighting of evidence in the decision^41^, as quantified by the analysis of psychophysical kernels^42–45^.

We, therefore, computed psychophysical kernels to determine the impact of the pharmacological manipulations on within-trial evidence weighting (Figure 3). We again used logistic regression, but now regressing the stimulus fluctuations around the generative mean at each position in the sequence on choice (Methods). At all positions did evidence fluctuations have a significant impact on the choice, demonstrating that participants tended to base their choices on the evidence integrated across samples. The shape of the psychophysical kernel revealed an overweight of earlier samples compared to later ones (primacy; Figure 3A), as previously observed^43,45^.

**Figure 3.**
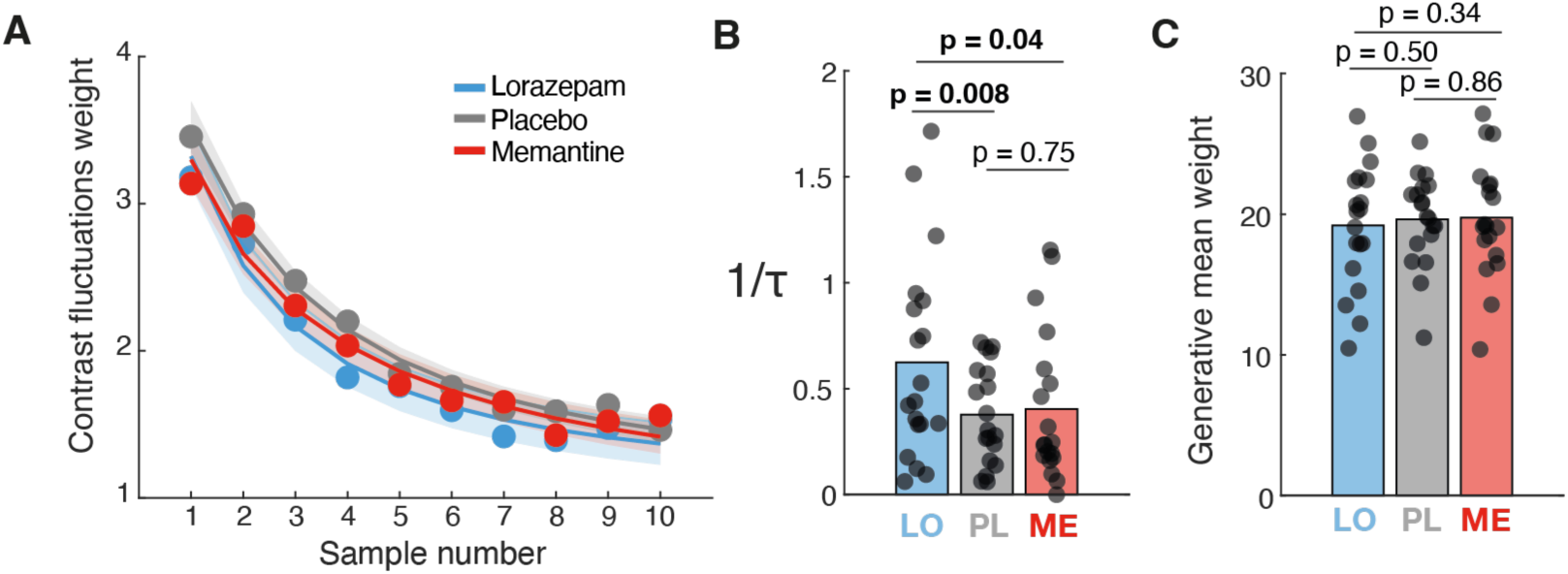
Timescale of evidence accumulation does not depend on memantine but on lorazepam. **(A)** Psychophysical kernel quantifying the impact of contrast difference fluctuations (residuals after subtracting the generative mean, Methods) on choice for different drug manipulations. Data points, group average. Lines, exponential decay function fits. Shaded area, SEM. **(B)** Exponential decay time constant for different drug manipulations obtained by fitting an exponential decay function to individual psychophysical kernels. **(C)** Impact of generative mean on choice for different drug manipulations. P-values are from two-sided permutation tests. Bars, group average; data points, individual participants.

We fitted an exponential decay function (see Methods) to the individual psychophysical kernels and estimated the decay time constant for each participant and drug condition. NMDA blockade via memantine did not have any detectable effect on the kernel shape compared to placebo. By contrast, the GABA-A boost via lorazepam shortened the decay time constant compared to both other conditions (Figure 3B). Neither drug had any effect on the sensitivity for overall evidence strength (Figure 3C), in line with our previous analyses of the psychometric functions (Figure 2A, C).

## Discussion

The NMDA receptor has been implicated in key cognitive capacities such as working memory^46,47^, evidence accumulation for decisions^20,34,41^, as well as learning and plasticity^48–50^ (but see^51^ for alternative models). Moreover, NMDA hypofunction has been implicated in cognitive disorders such as schizophrenia^24,52–54^. Here, we discovered a key contribution of NMDA receptors to history-dependent, adaptive biases in perceptual decisions under uncertainty.

Our results indicate that NMDA receptors shape the prominent sequential structure of human behavior. This is consistent with the hypothesis the NMDA receptors mediate short-term synaptic potentiation that retains information across trials in perceptual decision-making tasks. Support for this hypothesis has been derived from modeling behavior in the context of a continuous visuo-spatial working memory task in patients with (putative) NMDA hypofunction, such as individuals diagnosed with schizophrenia or anti-NMDA receptor encephalitis^24^, which may go along with other circuit alterations. Here, we directly and reversibly manipulated NMDA receptor function in healthy individuals in a placebo-controlled cross-over design. We studied adaptive choice history biases emerging in the context of perceptual categorizations performed in environments with different sequential correlation structures. It will be relevant in the future to test the effects of same pharmacological manipulations in healthy individuals performing the tasks used in the above clinical work, as well as the effects of the same clinical disorders on our perceptual categorization task.

The idea that NMDA receptors are a key synaptic feature for statistical learning and predictive processing has emerged from studies of different contexts. Mismatch negativity (MMN), a neural signature of predictive error, has been shown to be disrupted by the pharmacological NMDA receptor blockage in humans^55^ and animals^56^. Reduction in mismatch negativity^55,57^, serial dependence^24^ and choice history biases^28^ have been reported in patients with schizophrenia and psychosis-prone individuals, indicating a possible NMDA receptor dysfunction in these clinical populations. While the role of NMDA receptors in integrating information at timescales beyond single-trial processing in humans has so far been inferred mostly indirectly from clinical populations with NMDA receptor deficits, the present study provides direct support for this hypothesis in a healthy population by systematically modulating NMDA receptor function pharmacologically.

Our current approach also allowed us to identify dissociated effects of NMDA and GABA-A receptors on accumulation processes at different timescales that conspire to shape perceptual decisions: the accumulation of rapidly fluctuating sensory evidence within trials (on the order of hundreds of milliseconds) and the accumulation of experimental events (observations, actions, or outcomes) across timescales of up to tens of seconds. At the across-trial level, we found that participants were able to adjust their history biases to the statistical structure of the environment presented, as previously reported^7,16^, in a way that was well captured by an across-trial timescale accumulator model. At the within-trial level, subjects integrated the contrast fluctuations of all the ten input samples with a systematic overweight of earlier samples compared to later ones.

The lack of a memantine effect on stimulus evidence accumulation in our current task does not imply that the NMDA receptor plays no role in this process. Indeed, necessity of that receptor for integrating rapidly fluctuating evidence is postulated by current circuit models of perceptual decision-making. In these models, the necessity results from the long synaptic timescale of NMDA receptors and, more importantly, its predominance on recurrent cortical connections necessary for network reverberation^34,38,41^. It is possible that memantine (or other NMDA receptor blockers, such as ketamine) alter evidence weighting profiles when administered in a higher dosage. In line with this interpretation, a recent study found an enhanced effect of disambiguating sensory information on the perception of ambiguous stimuli under systemic NMDA blockade using ketamine infusion in humans^58^. However, ketamine did not evidently alter the evidence accumulation profile, but instead affected participants’ propensity to make premature decisions^59^. Another possibility is that the lack of effect of memantine on evidence accumulation results from the combined effects of that blockade of NMDA receptors on both, excitatory and inhibitory cells, which may cancel. Previous circuit modeling work has manipulated the NMDA conductance only on one of these connection types^41,52^. Future work could test these possible scenarios using a combination of modeling and NMDA blockade restricted to specific cortical area, similar to what was done for working memory^46^.

The effect of lorazepam found in the current study appears to be inconsistent with modeling work^41^ that predicts a flattening of the evidence accumulation profile under increased inhibition. However, recent physiology work indicates that the primacy profiles in psychophysical kernels reflect not just of evidence accumulation mechanisms in downstream decision circuits, but also an attenuation in the encoding of the evidence in sensory cortex^43,60^. In a variant of our contrast averaging task, primary visual cortex (V1) encodes individual contrast samples with a precision that decays over time in a manner that contributes to the primacy evident in the psychophysical kernels estimated from behavior^43^. GABAergic signaling from inhibitory interneurons plays a key role in sensory adaptation^61^. This mechanism may account for the lorazepam-induced effect on evidence weighting profile we observed here.

Recent work reported that pharmacological boost of catecholamine levels in humans through atomoxetine also reduces history biases, in a similar perceptual decision-making task^62^. This effect may reflect a down-weighting of the impact of recent history through catecholamines, in particular noradrenaline, as proposed by an influential theoretical framework^63^. How this effect of noradrenaline is mediated at the cellular and synaptic level is so far elusive. One possibility in line with our current finding is that noradrenaline reduces NMDA-R dependent synaptic plasticity mechanisms, which incorporate the build-up of history-dependent prior expectations.

History-dependent biases may sometimes, but not always, be reflected in biased persistent cortical activity between trials. Some neurophysiological studies have observed such persistent trial history codes in posterior parietal^10,64–66^, prefrontal^23,67^ and motor cortex^68^. Specifically, motor cortical activity tends to carry a trace of previous actions that biases the subsequent action (typically toward alternation)^68^. The build-up of action-selective motor cortical activity may also be shaped by history bias, without any clear history trace in the pre-trial baseline activity^16^. History bias may be encoded in an action-independent format in prefrontal^23^ and parietal^65,69,70^ persistent activity (see ^10^ for action-coding persistent parietal activity). Yet, also in these association areas, other studies found little evidence for such stimulus-selective persistent activity between successive trials, and instead linked history biases to activity-silent mechanisms, where synaptic efficacies carry information over trials^23,24,72^. Overall, the involvement of persistent cortical activity in history-dependent biases seems to be highly context-dependent. The involvement of synaptic traces mediated by NMDA receptors may be a more consistent mechanism.

By showing a direct effect of NMDA receptors on history biases in healthy individuals, these results provide a mechanistic link between synaptic function and perceptual decisions formation, offering possible new insights into the neurobiological mechanisms of clinical conditions such as schizophrenia.

## Methods

### Participants

Twenty-three healthy human participants (mean age 28, range 21-40, 9 females, 19 right-handed) took part in the study after informed consent and introduction to the experimental procedure. The study was approved by the ethics review board of the Hamburg Medical Association responsible for the University Medical Center Hamburg-Eppendorf (UKE). Exclusion criteria, all of which were assessed by self-report, were: history of any neurological or psychiatry disorders, hearing disorder, history of any liver or kidney disease or metabolic impairment, history of any chronic respiratory disease (e.g., asthma), history of hyperthyroidism or hypothyroidism, pheochromocytoma (present or in history), allergy to medication, known hypersensitivity to memantine or lorazepam, family history of epilepsy (first or second degree relatives), family history of psychiatric disorders (first or second degree relatives), established or potential pregnancy, claustrophobia, implanted medical devices (e.g., pacemaker, insulin pump, aneurysm clip, electrical stimulator for nerves or brain, intra-cardiac lines), any non-removable metal accessories on or inside the body, having impaired temperature sensation and / or increased sensitivity to heat Implants, foreign objects and metal in and around the body that are not MRI compatible, refusal to receive information about accidental findings in structural MR images, hemophilia, frequent and severe headaches, dizziness, or fainting, regularly taking medication or have taken medication within the past 2 months.

Three participants were excluded from the analyses: one due to excessive MEG metal artifacts and the other two due to not completing all six recording sessions. Thus, we report the results from n = 20 participants (7 females).

Participants were remunerated with 15 Euros for the behavioral training session, 100 Euros for each MEG session, 150 Euros for completing all six sessions, and a variable bonus, the amount of which depended on task performance across all three sessions. The maximum bonus was 150 Euros.

### Experimental design

The reported data come from a large pharmacological intervention study using a double-blind, randomized, crossover experimental design. Each participant completed six experimental sessions that also entailing also magnetoencephalography (MEG) recordings during resting-state measurements as well as a simple auditory task described in previous work^30,71^. Each experimental session began with the oral administration of the NMDA receptor antagonist memantine^73^, the GABA-A receptor agonist lorazepam^74^, or a mannitol-aerosil placebo at low dosage (both drugs and the placebo each on two randomly selected sessions). This was followed by a 2.5 h waiting time, during which participants were kept under observation, and their blood pressure and heart rate was recorded every 15 minutes. The waiting period was chosen to jointly maximize plasma concentrations of both drugs during MEG recordings (spanning about 2 h): peak plasma concentrations are reached ∼3 to 8 h after memantine administration^75^ and 2 - 3 h after lorazepam administration^76^. Following the waiting period, participants were seated on a chair inside a magnetically shielded chamber and the experimental data collection commenced, including about 90 min of performance of the behavioral task used in the current study. This task around 3 h after drug intake.

We used the following dosages: 15 mg for memantine (clinical steady-state dose for adults: 20 mg) and 1 mg for lorazepam (common clinical daily dose between 0.5 and 2 mg). The substances were encapsulated identically to render them visually indistinguishable. The first and senior author are both certified medical doctors, and emergency medical care by the University Medical Center Hamburg-Eppendorf was available on campus at any time. The six sessions were scheduled at least one week apart to allow plasma levels to return to baseline (plasma half-life of memantine: ∼60 to 70 h^75^; half-life of lorazepam: ∼13 h^76^).

### Stimulus and behavioral task

Each trial of the task consisted of the following sequence of events (Figure 1A) After a variable delay (uniform between 0.5 and 1 s) ten successive samples of variable contrasts (100 ms each) were shown in each visual hemifield. Participants were asked to compare the average mean contrast across the left and right samples (i.e., forced choice report: “left is stronger” or “right is stronger”). The offset of the last sample marked the beginning of the response period for participants. Participants reported their binary choice (left vs. right), and their confidence about the correctness of that choice simultaneously, by pressing one of four different buttons, whereby the two hands were always mapped to different choices. The index and ring fingers of each hand were then used to report confidence. During MEG sessions, participants used two response pads, one for each hand. During the training sessions participants, used the same stimulus-response mapping, but pressed keys on a computer keyboard. After a participant’s response and a consecutive variable delay between 0 and 0.5 s auditory feedback was given (250 ms duration). A low tone indicated a wrong answer, and a high tone indicated a correct answer. The ten consecutive contrast samples from the two hemifields were drawn from two normal generative distributions of contrast levels whose mean difference (right - left) was centered on each participant’s 75% accuracy contrast level. This threshold was determined by running a QUEST staircase continuously in the background. The standard deviation of each of the two normal distribution was 0.15. To avoid confusion, we only presented those contrast sequences sample whose actual difference in (sample) mean contrast matched the sign of the mean difference of the selected generative distributions. After each block of 120 trials participants could take a short self-timed break. After the third block, participants took a longer break lasting at least five minutes. Each participant completed six sessions, whereby each session consisted out of six blocks of 120 trials, lasting approximately 90 min. Among the six blocks, four of them were defined as “neutral”, when the side of stronger stimulus was chosen random on each trial, and two of them were defined as “repetitive”, where the previous stimulus category was more likely to be repeated (80% repetition probability). The first session was a training session that took place in a behavioral laboratory and was used to expose participants to the task and to calibrate their performance to 75% correct. The subsequent six sessions were experimental recording sessions that took place in the MEG laboratory and yielded the data analyzed in this paper.

### Data acquisition

#### Behavior and eye tracking

Stimuli were generated using Psychtoolbox-3 for MATLAB and were back-projected on a transparent screen using a PROPixx projector at 120 Hz during MEG recordings, or on a VIEW-Pixx monitor during the training session in a behavioral psychophysics laboratory. Eye movements and pupil diameter were recorded at 1000 Hz with an EyeLink 1000 Long Range Mount system (equipment and software, SR Research).

#### MEG

We used a CTF MEG system with 275 axial gradiometer sensors and recorded at 1200 Hz, with a (hardware) anti-aliasing low-pass filter (cutoff: 300 Hz). Recordings took place in a dimly lit magnetically shielded room. We concurrently collected eye-position data with a SR-Research EyeLink 1000 eye-tracker (1000 Hz). We continuously monitored head position by using three fiducial coils. After seating the participant in the MEG chair, we created and stored a template head position. At the beginning of each following session and after each block we guided participants back into this template position. We used Ag/AgCl electrodes to measure ECG and vertical and horizontal EOG.

#### Magnetic resonance imaging

Structural MRIs were obtained for each participant. We collected T1-weighted magnetization prepared gradient-echo images (TR = 2300 ms, TE = 2.98 ms, FoV = 256 mm, 1 mm slice thickness, TI = 1100 ms, 9° flip angle) with 1 × 1 × 1 mm^3^ voxel resolution on a 3 T Siemens Magnetom Trio MRI scanner (Siemens Medical Systems, Erlangen, Germany). Fiducials (nasion, left and right intra-aural point) were marked on the MRI.

### Behavioral data analysis

Behavioral data were analyzed with customized scripts (see associated code, which will be made available upon publication).

#### Logistic regression model with history bias

We quantified the influence of the history of previous choices and stimulus categories on the current choice, by using a logistic regression model with a history-dependent bias term that shifted the psychometric function along the horizontal axis:

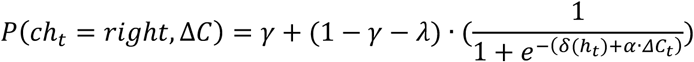

Where γ and λ are the probabilities of stimulus-independent errors (i.e., lapses), Δ*C_t_* is the delta contrast (right-left) at trial t, α is contrast sensitivity and δ(*h_t_*)is a bias term:

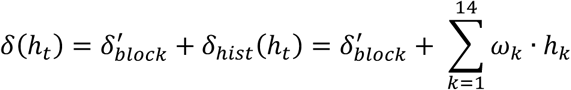

that is, the offset of the psychometric function, consisted of an overall block-specific bias δ^1^ for one specific choice and a history-dependent bias term δ_hist_(*h*_t_) = ∑_k=1_^14^ ω_k_ · *h*_k_, which was the sum of the preceding 7 choices *c*_t-1_ to *c*_t-7_and the preceding 7 stimulus categories *z*_t-1_ to *z*_t-7_, each multiplied with a weighting factor ω_k_. Right and Left choices and stimulus categories were coded as 1 and –1. The weighting factors ω_k_ specified the influence of each of the n preceding choices and stimulus categories on the current choice. Positive values of ω_k_ referred to a tendency to repeat, and negative values of ω_k_ referred to a tendency to alternate the choice or stimulus category at the corresponding lag. The parameters were estimated by minimizing the negative log-likelihood using MATLAB *fmincon* function. A regularization penalty on the lapse parameters γ and λ was added to the negative log-likelihood function so that:

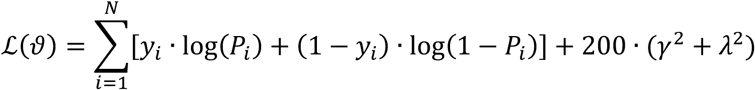

Where *y*_i_ ∈ {0,1} represented the observed choice on trial i, *P*_i_ is the probability of a right choice given by our logistic function and parameters γ and λ are lapse rates.

#### Psychometric curves of choice repetition

We quantified how the probability of repeating the previous stimulus side changed with the repeating stimulus evidence for the two environments. The repeating stimulus evidence *e*_t_ was defined by multiplying the current delta contrast Δ*C* with the previous trial stimulus side (*s*_t-1_ = {−1,1}). Psychometric curves were fitted to a probit function:

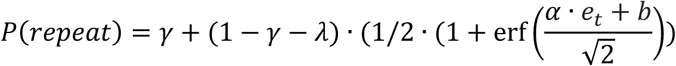

Where *e*_t_ is repeating stimulus evidence, α quantifies the discrimination ability, γ and λ are the probabilities of stimulus-independent errors (i.e., lapses), and the repeating bias *b* captured participants’ tendency to repeat (*b* > 0) or alternate (*b* < 0) their previous stimulus side. A regularization penalty on the lapse parameters γ and λ was added to the negative log-likelihood function so that:

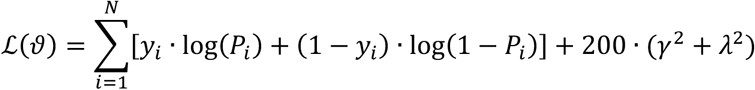

Where *y*_2_ ∈ {0,1} represented the observed choice on trial i, *P*_2_ is the probability repeat previous stimulus side given by our probit function and parameters γ and λ are lapse rates.

### Generative model of history bias

#### Model description

We adapted the dynamical model developed by^12^ to quantify how much previous choices, stimulus categories and transitions affected the current choice. The model contained three latent variables zL, zT, zS that were updated at each trial t.

*zS* represents the current stimulus:

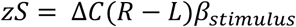

Where Δ*C*(*R* − *L*) is the average delta contrats (right-left) across the ten samples, and α quantifies contrast sensitivity.

*zL* represents the lateral module and was updated as:

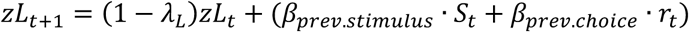

Where λ_<_ is a leak parameter (from 0 to 1), β_prevstumus_ and β_prev.choice_ are the update parameters for previous stimulus category *S*_t_ and previous choice *r*_t_ respectively.

*zT* represents the transition module and was updated as:

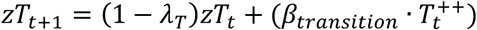

Where λ_B_ is a leak parameter (from 0 to 1), β_prev.stumus_ is the update parameters for previous transitions where both last two trials outcomes were correct choices (*T*^t^).

The accumulated transition evidence *zT* maintained a running estimate of the transition statistics and its transformation onto the transition bias γ_B_ was gated by second variable G by setting:

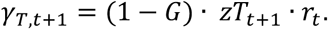

The parameter G was bounded between [0,2], where G=1 indicates a reset of transition bias after error trials, G=2 indicates that the transition bias was reversed after errors, and G=0 indicates that the transition bias is not gated. The gate parameter *G* was set to 0 after correct trials.

The probability of right response was then defined by:

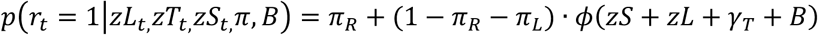

Where π_R_ and π_L_ are right and left lapses, *B* is a fixed side bias, and φ is a probit function.

#### Fitting procedure

We defined a Gaussian prior with zero-mean diagonal covariance for all model parameters except lapse parameters, and a Dirichlet distribution for the lapse parameter π:

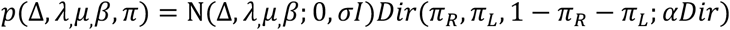

We used σ = 1 and α Dir = (1; 1; 38) (yielding an overall prior lapse probability of 5%).

Parameters in the latent model (both with and without modulated biases) were fitted by maximizing the log-posterior of the observed responses (i.e. the sum of the log-prior and the log-likelihood). Maximization was performed using function NLopt in R, using BOBYQA algorithm^77^.We run the minimization procedure with 500 randomly selected initial points.

#### Psychophysical kernels

Psychophysical kernels were computed using the logistic regression model:

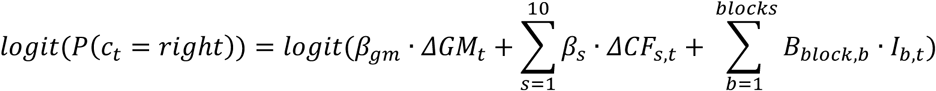

Where Δ*GM*_t_ is the delta generative mean (right-left) from the right and left generative distributions of sample contrast at trial t, Δ*CF*_st_ are the delta contrast fluctuations around the delta generative mean (right-left) across the 10 samples *s*, and *I_b,n_* is a binary indicator variable coding the block identity at trial t. The model was fit on trials from both repetitive and neutral blocks and psychophysical kernels were defined by the values of β_3_ across the ten samples.

To quantify the shape of the psychophysical kernels we fitted to the 10 β_3_ an exponential decaying function with an offset parameter:

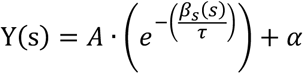

Where A controls the initial amplitude, τ is the decaying time constant and α is the offset parameter.

