## Supplemental Information for "History-dependent Biases in Perceptual Decisions Depend on NMDA Receptors"

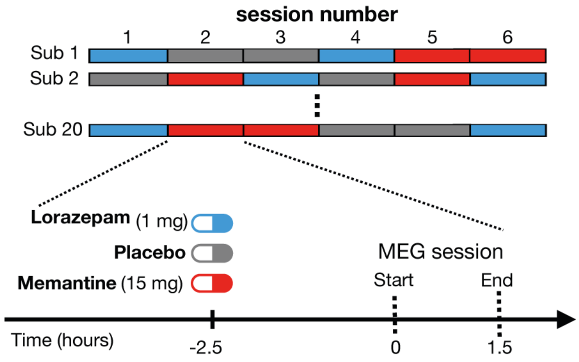


**Figure S1. Experimental design and drug manipulations**. Subjects underwent six MEG sessions following intake of placebo, lorazepam or memantine and 2.5 h waiting period.

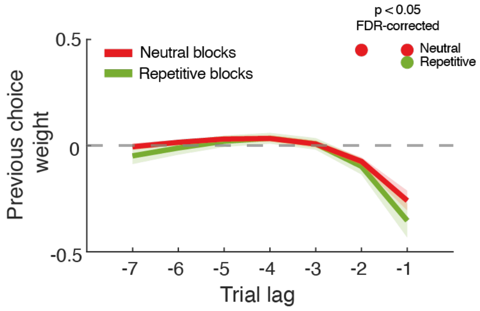


**Figure S2. Impact of previous choices on current choice.** Beta weights of previous choices as a function of trial lag for Repetitive and Random environments. Lines, group average, shaded area, SEM. Upper marks, $p < 0.05$; FDR-corrected**.**

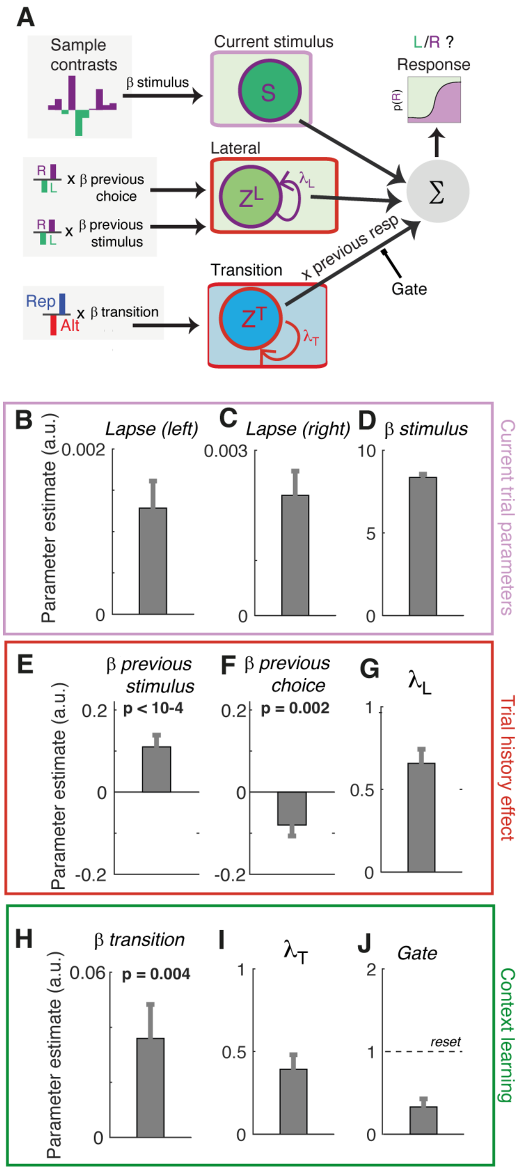


**Figure S3. Generative model of adaptive choice behavior across sensory environments.** (A) Schematics of the model. The sensory module accumulates perfectly (i.e. no leak) samples of delta contrast from the current stimulus. The lateral module accumulates lateral evidence ZL (i.e. evidence to choose right vs left), with a certain leak $\lambda_L$ and with inputs from the previous choice and previous stimulus category. This module recapitulates with just three parameters the previous stimulus category kernel (Fig. 1J) and the previous choice kernel (Fig. S2). The transition module accumulates the transition evidence ZT which differently from the later module represents evidence to repeat (ZT >0) or alternate (ZT<0) the previous response (Hermoso-Mendizabal et al 2020). The transition evidence has a leak $\lambda_T$and it is updated with the previous transition T: this variable is T = +1 if the two last choices were correct and to the same side (a correct repetition), T= -1 with the two last choices were correct and to different sides (a correct alternation) and T = 0 if at least one of the two choices was incorrect. The transition evidence recapitulates into a single variable the learning of the repeating statistics of each block: during a random block ZT would fluctuate around zero whereas in the repetitive block it will fluctuate around positive values. When looked under the GLM used in the main text, this transition evidence can generate the difference in previous stimulus kernel shown in Fig. 1J. The impact of the transition evidence onto the current choice is then obtained from the product of ZT and previous choice after correct trials. After error trials, we implemented an after-error modulatory parameter named Gate such that the impact Zt x (previous choice) x (1-Gate). This parameter could reduce the impact of ZT (0 < Gate < 1) or reverse it (1 < Gate < 2). (B-J) Best fitting values of model parameters. (B-D) Present trial parameters: stimulus independent lapses (B-C) and current stimulus impact $\beta_{stimulus}$ (D). (E-J) Trial history parameters: weight on previous stimulus categories $\beta_{previous\ stim}$ (E) and previous choices $\beta_{previous\ choice}$ (F) which are accumulated with a leak term $\lambda_L$ (G). weight on previous correct transitions $\beta_{transition}$ which are accumulated leak term $\lambda_T$ (I). Accumulated transition evidence is attenuated in the fitted model after error trials (J). P-values are from two-sided permutation tests against 0. Bars, group average; errorbars, SEM.

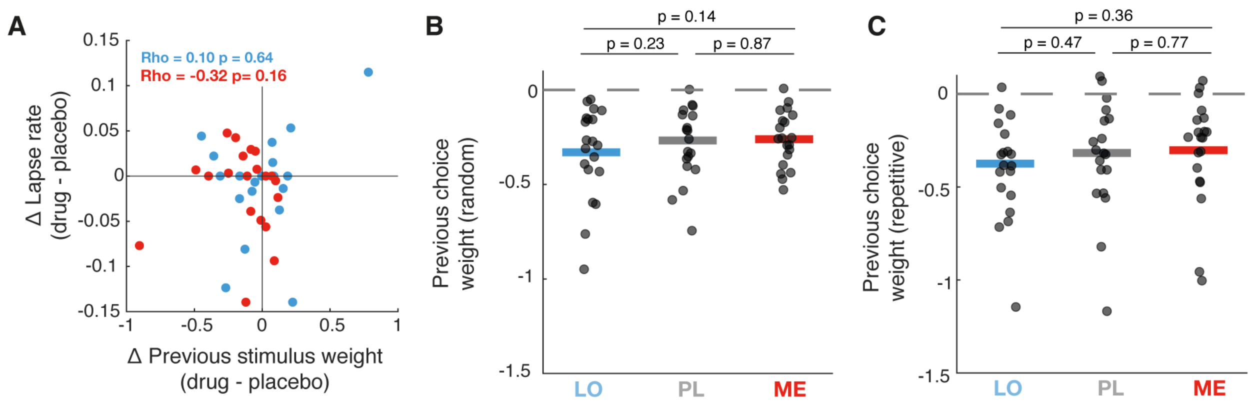


**Figure S4. Drug-induced changes in stimulus history bias. (A)** Correlation between drug-induced changes in previous stimulus weights obtained from behavioral model and lapse rates (Repetitive environment). Dots are participants. **(B-C)** Impact of previous choice in random (B) and repetitive (C) environment. P-values in all panels are from two-sided permutation tests. Horizontal lines, group average; data points, individual participants.
